# Microfluidic Chip for Label-Free Removal of Teratoma-forming Cells from Therapeutic Human Stem Cells

**DOI:** 10.1101/830588

**Authors:** Kyle Wellmerling, Christian Lehmann, Ankur Singh, Brian Kirby

## Abstract

Teratoma formation remains a safety concern in therapeutic cells derived from human-induced pluripotent stem cells (hiPSCs). Residual Teratoma forming cells are present in small numbers in differentiated hiPSC cultures and yet are of significant roadblock to the manufacturing and clinical translation of stem cell therapies. Rare cells are often difficult to remove with standard flow cytometry or magnetic bead sorting techniques. Here, we first characterized time-dependent expression of a teratoma marker, stage-specific embryonic antigen (SSEA)-5, which binds the H type-1 glycan during neural differentiation of hiPSCs. We engineered a microfluidic geometrically enhanced differential immunocapture (GEDI) technology to remove SSEA-5+ rare cells from hiPSC-derived neural progenitor cells (hiPSC-NPCs). The GEDI chip presents a facile tool to potentially functionalize with multiple antibodies and robustly enhance hiPSC-derived cell population safety prior to therapeutic transplantation. The approach is potentially amenable to generate a wide variety of high-quality therapeutic cells and can be integrated within the pipeline of cell manufacturing to improve patient safety and reduce the cost of manufacturing through early removal of undesirable cell types.

## 1 Introduction

Pluripotent stem cells (PSCs) represent a highly promising strategy to derive cells for therapeutic transplantation[1–5]. Reprogramming of somatic cells to human-induced PSCs (hiPSCs)[6] has emerged as a powerful technology for modeling diseases and as a method of generating patient-specific cells for therapeutic purposes. Differentiated cells derived from patient-specific hiPSCs are thought to be less immunogenic than tissue derived from non-allogenic human embryonic stem cells (hESCs)[7–9], circumvent the ethics of using hESCs, and self-renew indefinitely, enabling derivation of the large number of cells required for therapeutic transplantation[10].

Of tissue derived from hiPSCs, neural stem and progenitor cells are a promising therapy for treating chronic neurodegenerative diseases, for example Alzheimer’s disease[11] and Parkinson’s disease[12, 13]. Neural progenitor cells (NPCs) are a promising strategy to treat Alzheimer’s disease and Parkinson’s disease due to a widespread loss of neurons being the prevailing consequence of this diseases[14–16]. Cell transplant therapies that enhance neurogenesis or replace lost neurons may delay the progression of neurodegenerative diseases. NPC grafts onto rodent models of AD demonstrate increased hippocampal synaptic density and increased cognitive function[17]. Additionally, cellular therapies that used fetal ventral midbrain tissue as a source of dopaminergic neurons have reported varying degrees of success[18]. However, cellular transplants utilizing fetal ventral midbrain tissue are limited due to ethical concerns of harvesting cells from donors and the inability to obtain adequate numbers of cells for transplant. hiPSC-derived NPCs (hiPSC-NPCs) are poised to overcome these obstacles and have demonstrated cell integration into host tissue[19]. In addition to chronic neurodegenerative diseases, acute neurodegeneration may result from traumatic brain injury (TBI) or spinal cord injury[20–22]. Human NPC transplantation has shown promising results to treat TBI and has demonstrated neuroprotection and improved cognitive outcomes following TBI[23]. Taken together, these results demonstrate that NPCs are a promising cellular therapy to treat acute neurodegeneration in humans. However, despite the promise of hiPSCs derived cells to treat disease, tumorigenicity of residual pluripotent-like cells in differentiated stem cells remains a concern[24, 25]. At present, numerous techniques have been applied to remove teratoma-forming potential from hiPSC-derived products, such as pretreatment with antimitotic agents and delayed transplant[26], sorting with flourescent activated cell sorting (FACS)[27], magnetic activated cell sorting (MACS)[28], introduction of inducible suicide genes[29], and complete differentiation into mature dopaminergic neurons[30]. Unfortunately, each of these methods have limitations. Pretreatment with antimitotic agents and delayed transplant or complete differentiation into mature dopaminergic neurons increases turn-around time to derive therapeutic cells, and could make it difficult to transplant cells during the short optimal timing of transplantation in the subacute phase[31]. Although inducible suicide genes respond in 95% of cells, they are unlikely to eliminate teratoma because as few as 2 embryonic stem cells in two million non-neoplastic cells are capable of forming tumors in 60% of the transplants[26, 29]. Additionally, FACS and MACS have thus far proven insufficient to eliminate teratoma-forming potential and both require pre-labeling cells with antibodies, which could interfere with proliferation and differentiation[27, 28].

Microfluidic devices represent a potential avenue to remove rare, teratoma-forming cells. Microfluidic devices have shown effective capture of rare circulating tumor cells (CTCs) from whole blood[32], indicating their potential to remove rare cells from a cell population. Further, microfluidic devices have demonstrated enrichment and high survivability of hiPSCs and derived neural stem cells[3]. To this end we have engineered a geometrically enhanced differential immunocapture (GEDI) chip to capture and remove rare, teratoma-forming cells from a hiPSC-NPC population by exploiting the expression of stage-specific embryonic antigen (SSEA)-5, which strongly binds the H type-1 glycan and is specifically expressed on hPSCs. The GEDI chip exploits size of cells to maximize the number of collisions a cell experiences with an antibody functionalized post (**Figure 1**). Device geometry is designed such that cells above a set size follow a pathline through the device that causes repeated collision with the antibody functionalized posts. To remove cells from a hiPSC-NPC population, cells are first differentiated into NPCs, resuspended as single-cells, then teratoma-forming cells are captured by antibodies on the chip surface, and a purified population is obtained in the effluent (**Figure 1d**). We believe this microfluidic immunocapture represents a promising tool to address the tumorigenic cells, potentially reduce the cell manufacturing costs by reducing post-bioprocessing of scaled-up cells and enable translation of safer stem cell therapies.

**Figure 1.**
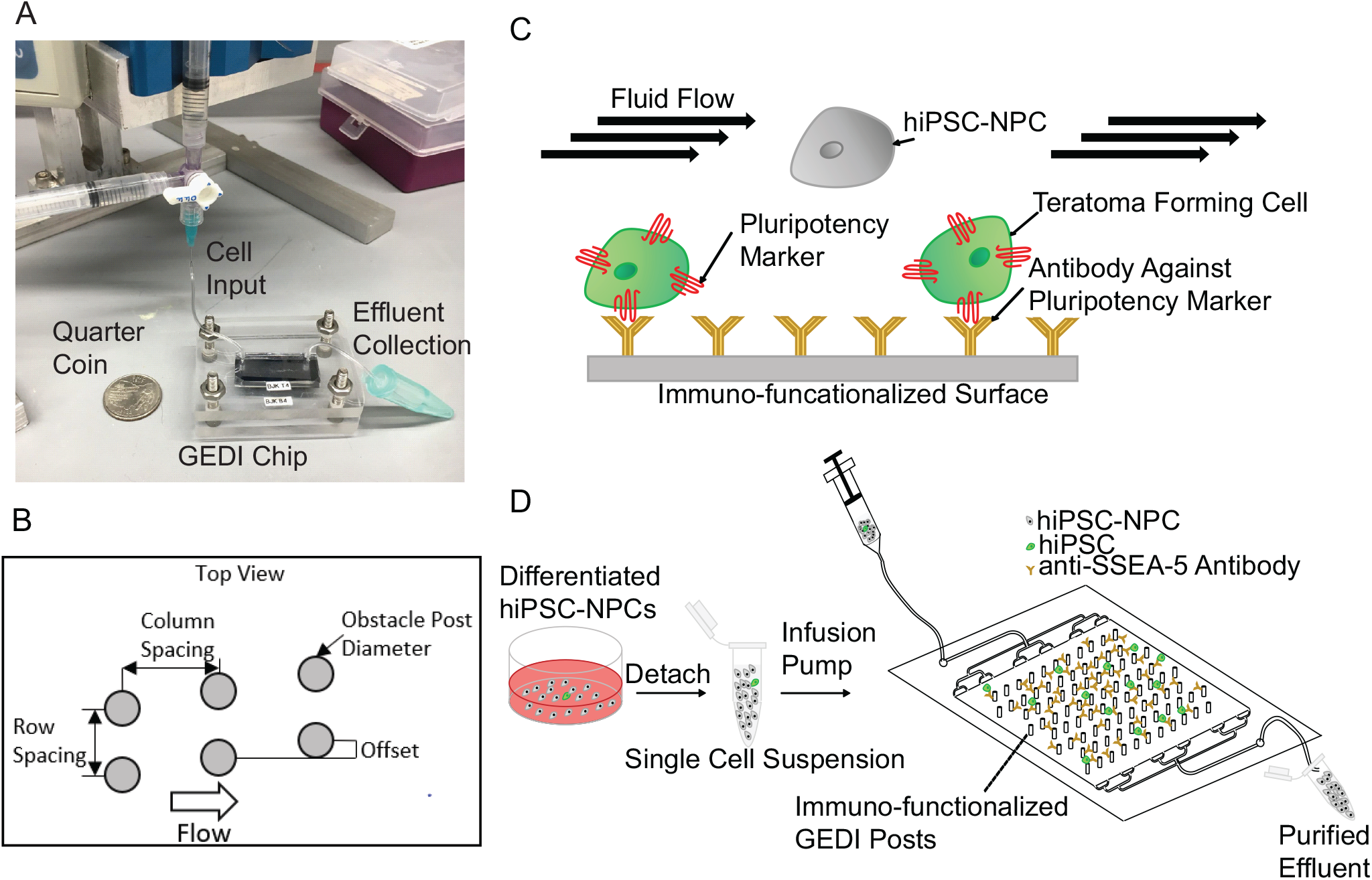
Schematic of GEDI chip and experimental procedure. (**A**) Picture of GEDI chip setup. (**B**) GEDI chip schematic (**C**) Cartoon depiction of surface functionalization. (**D**) Overview of GEDI experimental procedure.

## 2 Results and Discussion

### 2.1 Time-dependent expression of SSEA-5 on hiPSCs and hiPSC-NPCs

To design a device that removes teratoma-forming cells from hiPSC-NPCs we first sought to characterize the differentiation dynamics of hiPSCs. We sought to characterize how SSEA-5 expression changed during neural differentiation, as SSEA-5 was our principal marker for cells with high teratoma-forming potential[27] rather than cells solely expressing TRA-1-60. We used flow cytometry to measure single-cell SSEA-5 expression at different time points of hiPSC differentiation and measured the percent of the population SSEA-5+ at each time point of differentiation (**Figure 2a**). During differentiation we noted that SSEA-5 expression decreased dramatically throughout the 12-day differentiation period (**Figure 2b,c**).). Interestingly, SSEA-5 expression dropped at discrete intervals rather than continuously during the differentiation process. SSEA-5 expression was roughly constant during the first 3 days of neural differentiation and then dropped dramatically at day 4. SSEA-5 expression remained constant again through day 6 and dropped further between day 6 and day 9. After 9 days, SSEA-5 expression levelled off, indicating that most cells have completed differentiation by day 9.

**Figure 2.**
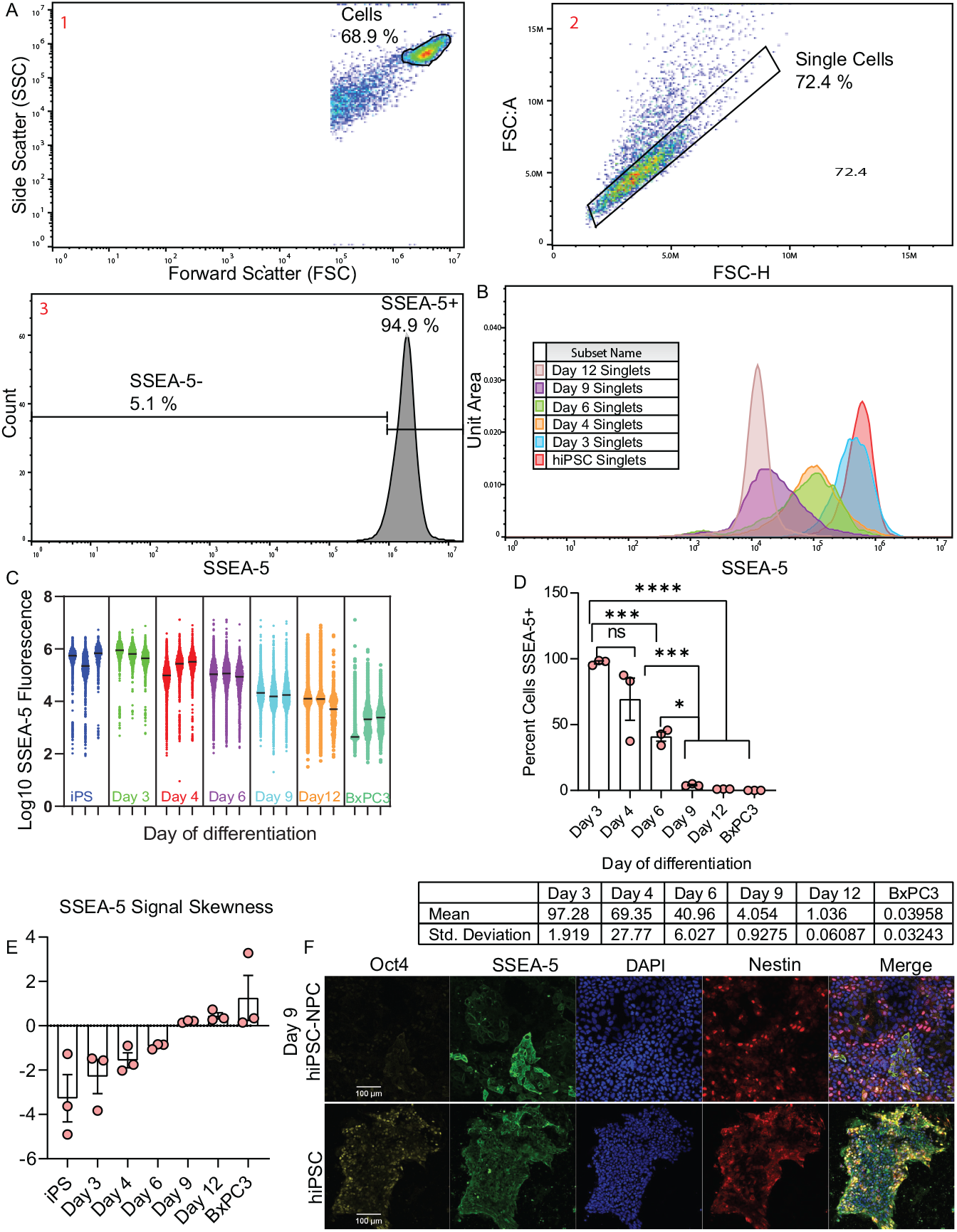
Temporal expression of SSEA-5 on hiPSCs and hiPSC-NPCs. (**A**) Gating of single-cells from SSC:H vs. FSC:H to identify cells, FSC:A vs FSC:H to identify single-cells, and thresholding of hiPSCs to identify SSEA-5+ cells (averaged over 3 hiPSCs populations. (**B**) SSEA-5 expression histograms for different time points of differentiation. (**C**) Log10 SSEA-5 expression for n=3 biological replicates at each time point. (**D**) Percent of cells SSEA-5^+^ at each time point (bar represents mean), *p=.05, ***p<.001, ****p<.0001. (**E**) Skew of SSEA-5 expression at each time point (error bars represent mean with SEM). (**F**) Immunofluorescence Images of day 9 hiPSC-NPCs (top) and undifferentiated hiPSCs (bottom) (10x)(scale bar: 100*μm*).

However, despite the drop in average SSEA-5 expression, we sought to investigate whether there existed a potential residual population of pluripotent or partially-differentiated cells with teratoma-forming potential. For each differentiation time point, we observed the percent of cell population with SSEA-5 expression above the bottom 5% of undifferentiated hiPSCs. We designated this fraction as SSEA-5+ (**Figure 2d**). Strikingly, we note that on day 12 of differentiation, 1 ±.02% of the cells were still SSEA-5+ after a full time-course of differentiation. It is unlikely this small fraction of cells represents noise in the flow cytometry because control non-stem cells (BxPC3 cell line) stained for SSEA-5 only showed 0.04 ±.02% of the population positive for SSEA-5. Similar results were obtained with the use of a single antibody conjugated to a fluorophore to measure SSEA-5 expression (**Supplementary Figure S1a**). This suggests that the observed differences were not measurement errors but indeed there is a residual population of hiPSC-NPCs with a pluripotency signature and thus teratoma-forming potential. To further investigate these cell populations we plotted the skew of SSEA-5 expression at each time point of differentiation. Interestingly, the differentiating cells were initially skewed towards low SSEA-5 expression (negative skew), indicating a few cells lose pluripotency faster than the rest of the population. Later on, however, at day 9 and day 12 (**Figure 2e**), the skew became positive (skewed towards high SSEA-5). This indicates that at the later stages of differentiation there are cells that slowly differentiate and retain pluripotency expression. Finally, to corroborate flow cytometry data with immunofluorescence (IF) images (**Figure 2f**), we stained undifferentiated hiPSCs and day 9 hiPSC-NPCs with Oct4, a master regulator of pluripotency, SSEA-5, and Nestin (a marker of neural differentiation). Strikingly, IF images corroborate flow data, clearly indicating that pockets of cells with high SSEA-5 expression remain on day 9, though with weak Oct4 expression, suggesting the cells retain partial pluripotency.

### 2.2 Microfluidic immunocapture of teratoma-forming cells

Having confirmed the presence of residual cells with expression of pluripotency markers and thus teratoma-forming potential, we sought to employ a microfluidic GEDI chip to remove the teratoma-forming cells from hiPSC-NPCs. To demonstrate SSEA-5 as a pluripotency surface marker that is capable of capturing hiPSCs on a GEDI chip, we first qualitatively compared capture on glass coverslips functionalized with anti-SSEA-5, anti-TRA-1-60, and normal mouse IgG antibodies (isotype control) (to test for steric hindrance (**Figure 3a**, as reported earlier by us[33]). We noticed similar numbers of cells on the anti-SSEA-5 and anti-TRA-1-60 coverslips compared to the isotype control functionalized cover slips. Knowing that anti-SSEA-5 and anti-TRA-1-60 coverslips can capture hiPSC, we tested functionalized GEDI chips with each of the above mentioned antibodies, either alone or in combination of anti-SSEA-5 and anti-TRA-1-60. Undifferentiated hiPSCs were flowed through the GEDI chips using a syringe pump and at concentrations of approximately 300 cells/mL. A pass through rate was then calculated based on the number of cells counted in the effluent of each chip (ratio of number of cells in effluent to number of loaded cells). Observed pass through rates are indicated in **Figure 3b**. Whereas anti-SSEA-5 and anti-TRA-1-60 GEDI chips had similar pass through rates, the isotype-control GEDI chips had a much higher pass through rate, indicating both anti-SSEA-5 and anti-TRA-1-60 GEDI chips captured hiPSCs preferentially compared to the control. Given that the anti-SSEA-5 GEDI chips had the lowest overall pass through rate, we chose to focus on this antibody for the remainder of studies in this paper.

**Figure 3.**
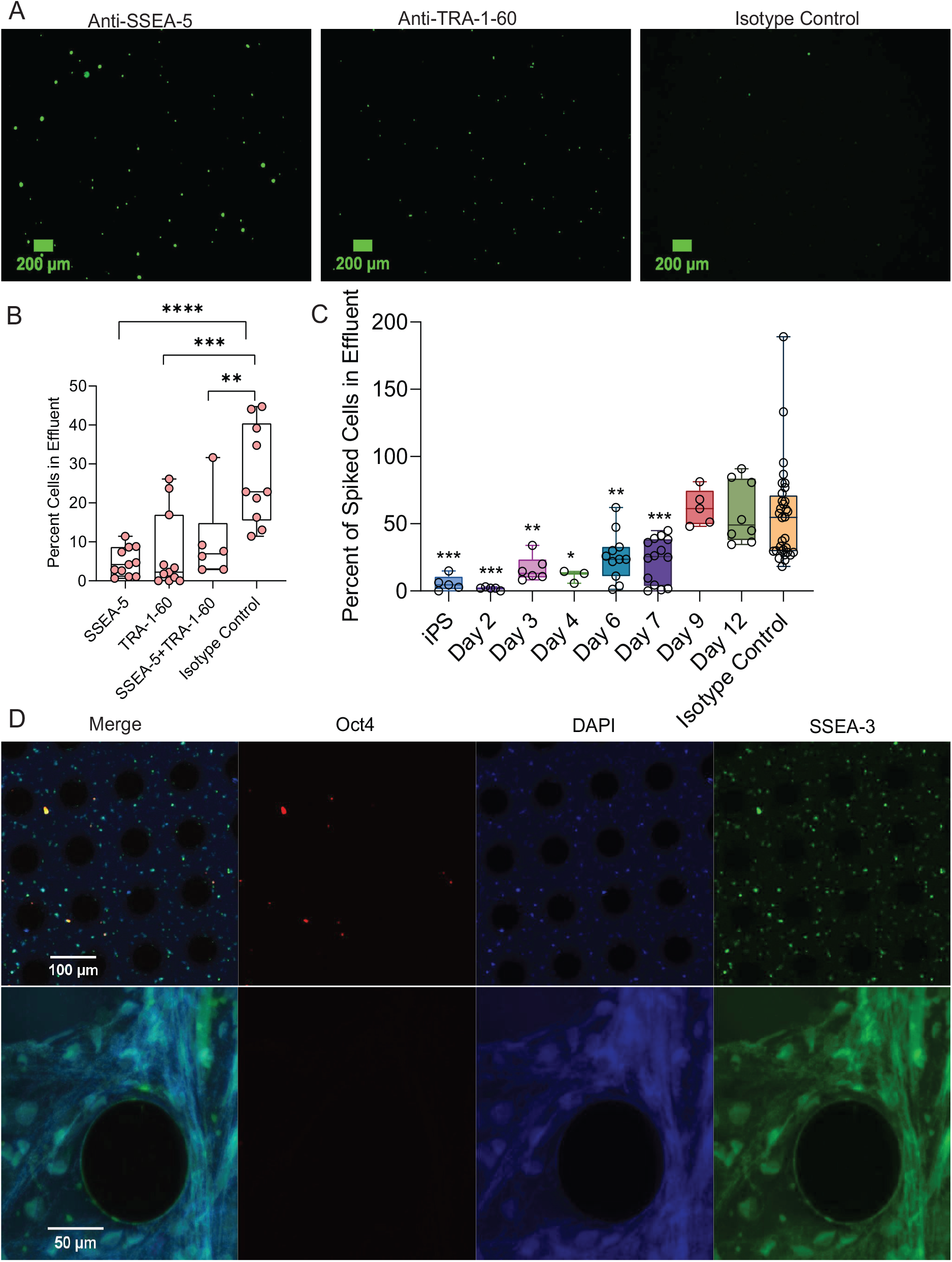
GEDI-enabled capture of teratoma-forming cells from undifferentiated hiPSCs and NPCs spiked into BxPC3s. (**A**) hiPSCs captured on antibody functionalized glass cover-slips. (**B**) Pass through rates for undifferentiated hiPSCs alone in suspension ****p<0.0001, ***p<0.001, **p<0.01, Tukey’s multiple comparison test (n=11 for SSEA-5, TRA-1-60, n=6 SSEA-5+TRA-1-60, n=10 isotype control). (**C**) Pass through rates for hiPSC-NPCs spiked into a larger BxPC3 population at different time points of differentiaiton (n=5 hiPSCs, Day 2, Day 9, n=6, Day 3, n=3, Day 4, n=11, Day 6, n=14 Day 7, n=8, Day 12, n=32 Isotype Control). (**D**) Immunofluorescence images of cells captured on anti-SSEA-5 GEDI chips. Top: 10x, Day 12 NPCs; Bot: 40x, P2 NPCs (hiPSC differentiated through 2 passages as hiPSC-NPCs, then proliferated through 2 passages in a neural proliferation media.

### 2.3 Capture based on SSEA-5 expression

After confirming SSEA-5 expression exists in differentiated hiPSC-NPCs, we sought to confirm our device’s ability to remove SSEA-5+ cells from artificially mixed cell population containing SSEA-5+ and SSEA-5-cells, in a manner proportional to SSEA-5 expression level of the target population. To demonstrate this capability, we spiked hiPSC-NPCs (300 cells/mL) with a population of 300,000 BxPC3 control cells/mL. We then calculated the pass through rate for each day of differentiation (**Figure 3c**), anticipating that with differentiation the number of SSEA-5+ cells will reduce and less hiPSC-NPC will be captured on the microfluidic pillars. Indeed, we found that as the hiPSC-NPCs became further differentiated, our removal specificity reduced because there were few SSEA-5+ cells to bind to SSEA-5 antibodies on GEDI pillars (cells in effluent for day 9 and 12 were similar to isotype control). This suggested that the GEDI chip efficiently capture based on SSEA-5 expression.

To confirm the cells that were captured on the GEDI chip were pluripotent stem cells with teratoma-forming potential, we processed day 12 hiPSC-NPCs through an anti-SSEA-5 GEDI chip and also tested purified NPCs that were maintained in proliferating cultures for 2 passages (P2 NPCs). We stained cells caught on the GEDI chip for Oct4, DAPI, and SSEA-3 (a stand in for SSEA-5, due to the inability to use a secondary fluorescent antibody against SSEA-5 on chip (**supplementary figure 1b**)) (**Figure 3d**). Strikingly, nearly all of the cells, irrespective of P2-NPC or hiPSC-NPC, caught on GEDI chip co-expressed DAPI and SSEA-3. However, in the day 12 hiPSC-NPC sample, a few Oct4+ cells were captured, which were not seen in the P2 sample, indicating further differentiation in the P2 sample, as expected.

### 2.4 Microfluidic Removal of SSEA-5+ Cells

Having confirmed that our device captures cells proportional to SSEA-5 expression and having confirmed the presence of rare teratoma-forming cells in differentiated populations, we quantified the effectiveness of the GEDI chip in removing teratoma- forming cells from a day 9 hiPSC-NPC population by imaging cells in the effluent of GEDI chips functionalized with anti-SSEA-5 and isotype control antibodies. The fraction of cells positive for pluripotency marker SSEA-3 was lower in the effluent obtained from anti-SSEA-5 GEDI chips than the isotype control chips for all thresholds (**Figure 4b**), indicating removal of pluripotent cells from the day 9 hiPSC-NPCs. Interestingly the fraction of OCT4+ cells was similar at low to moderate thresholds however was drastically lower for the SSEA-5 GEDI chip effluent at higher thresholds (**Figure 4c**). Further, the total fraction of Oct4+ cells was higher in day 9 hiPSC-NPCs compared to SSEA-3. A plausible explanation for the device performance is that, at low thresholds, bona fide NPCs are counted as OCT4+ cells, but at higher thresholds fewer bona fide NPCs are counted as positive, and thus residual pluripotent cells make up a larger fraction of the OCT4+ cells, of which there are more in the control GEDI chip effluent. Further, if we aggregate the average fluorescence intensity per nuclei area for the anti-SSEA-5 GEDI chips and isotype control GEDI chips and then normalize fluorescence intensity by the respective secondary control, we find reduced pluripotency marker expression in the anti-SSEA-5 GEDI chip compared with the isotype control chip (**Figure 4a**). This indicates the anti-SSEA-5 GEDI chips can sucessfully remove teratoma-forming cells from the day 9 hiPSC-NPC population and the effluent cells can then be potentially used for scale-up therapeutic purposes.

**Figure 4.**
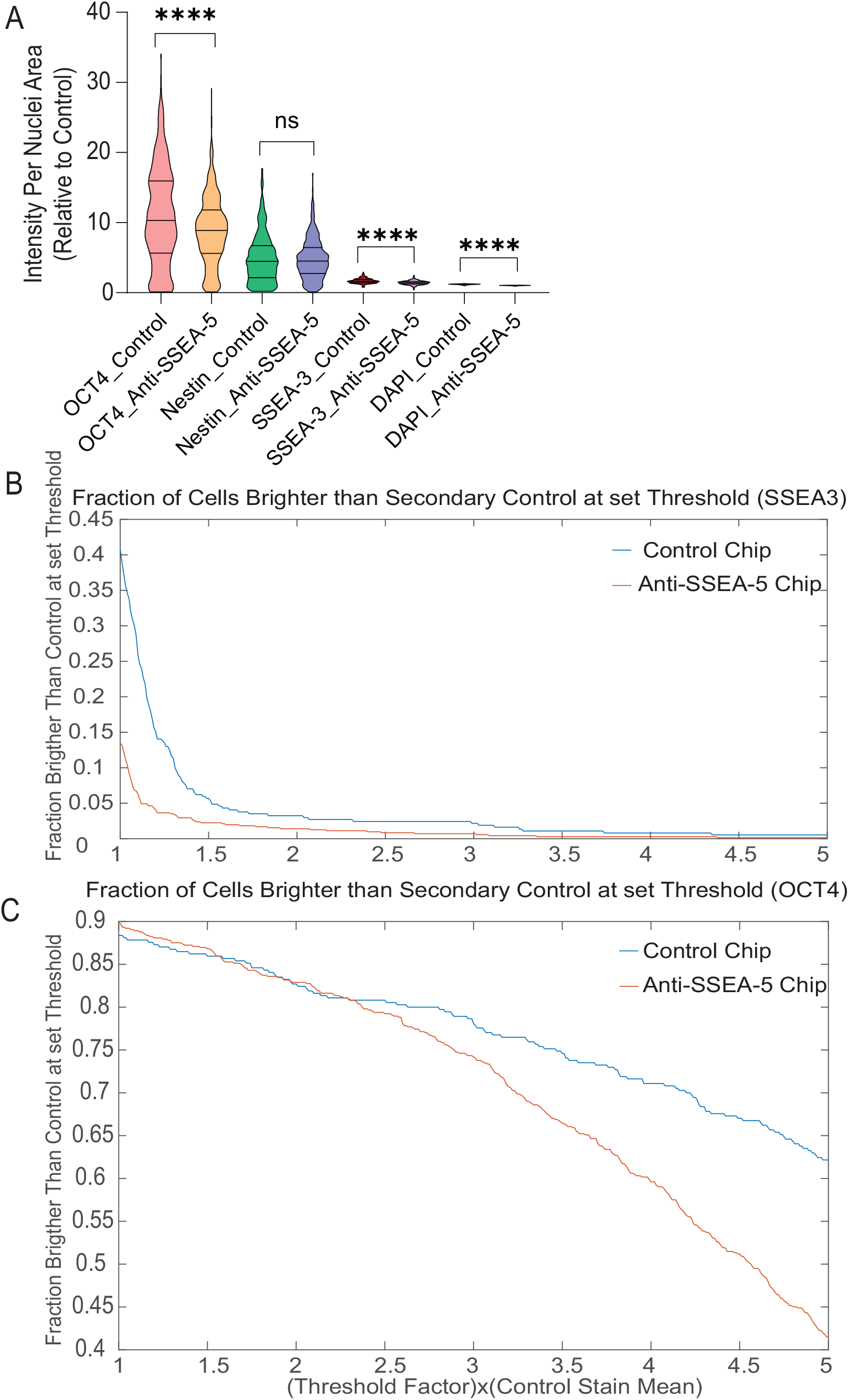
GEDI-enabled capture of residual teratoma-forming cells from hiPSC-NPCs. (**A**) Expression intensity per nuclei area relative to mean of secondary control for individual cells, ****p<0.0001, ns=not significant, unpaired parametric t test (n=370 cells for control, n=713 cells for anti-SSEA-5). (**B**) Fraction of day 9 hiPSC-NPCs (a) OCT4^+^ and (b) SSEA-3^+^ in effluent of anti-SSEA-5 and isotype control GEDI chip at varying threshold.

## 3 Conclusion

In summary, we have demonstrated that a small residual population of cells that express pluripotency markers and a specific glycan SSEA-5 exist even after prolonged periods of neural differentiation of hiPSCs. To this end, we have demonstrated a proof-of-concept study where a SSEA-5 antibody-functionalized GEDI chip can successfully reduce this teratoma-forming potential by filtering teratoma-forming cells out of the hiPSC-NPC population. We believe this work represents the first report on eliminating rare cells with teratoma-forming potential in hiPSC-derived neural stem cells with therapeutic potential. We believe that the GEDI technology is easily adaptable to other hiPSC-derived cells populations, could purify differentiated population in a label free manner, could be used to decrease turn-around time required to derive safe-therapeutic cell populations. Further we have demonstrated that hiPSCs lose SSEA-5 expression in a more punctuated or intermittent rather than continuous process, as indicated by drops in fluorescence intensity between day 3 and 6 cells and day 6 and day 9 cells. This suggests there may be optimal time-points at which to filter or sort cells prior to scale up for cell manufacturing and or therapeutic transplantation.

## 4 Methods

### 4.1 Device fabrication and functionalization

Devices were fabricated by A.M. Fitzgerald Associates (Burlingame, CA) according to previously described specifications[32, 34]. The silicon device surfaces were functionalized with a previously described protocol using 3-mercaptopropyltrimethoxysilane (MPTMS) and N-gamma-maleimidobutyryloxysuccinimide ester (GMBS) NeutrAvidin-biotin chemistry (Thermo Fisher Scientific, Rockford, IL)[32, 34].The devices were functionalized with primary antibodies via a biotinylation linkage. NeutrAvidin-functionalized devices were incubated with either (1) 2 *μ*g/mL biotinylated mouse anti-human anti-SSEA-5 antibody (Miltenyi Biotec, Bergisch Gladbach, Germany) (2) 10 *μ*g/mL mouse anti-human anti-TRA-1-60 antibody (ThermoFisher Scientific, Waltham, MA), (3) 2 *μ*g/mL of (1) and 10 *μ*g/mL of (2). Control devices were functionalized with biotinylated normal (non-specific) mouse IgG antibodies ((Miltenyi Biotec, Bergisch Gladbach, Germany) at 2 *μ*g/mL. All antibodies were prepared in 2% BSA with 1.3-1.5mM EDTA in PBS. Polydimethysiloxane (PDMS) sheets (5:1 base:curing agent), approximately 3 mm thick, were polymerized for 18 hr at 60°C and trimmed to form covers for the GEDI chip. A PDMS sheet was clamped to the top of the post arrays. Inlet and outlet holes were created with a biopsy punch, and tygon tubing was inserted into the PDMS gasket to connect GEDI chip inlet to a syringe pump and outlet to a microcentrifuge tube for effluent collection. Devices were primed with a 50/50 Ethanol/DI water mixture, and then flushed with DI water water and PBS before experiments.

### 4.2 Cell Culture

The iPSC cell lines were provided as a kind gift by the laboratory of Jan Lammerding, and were originally generated from healthy control subjects by Elisa Di Pasquale, as described previously[35, 36]. BxPC3 cells were obtained ATCC (Manassas, VA). All cell lines were cultured in humidified incubators (37 C and 5% CO2) using TeSR-E8 media (StemCell Technologies, Vancouver, Canada) or STEMdiff Neural Induction Medium (StemCell Technologies, Vancouver, Canada) for hiPSCs and hiPSC-NPCs respectively. hiPSC, hiPSC-NPCs, and NPCs were supplemented with 10 *μ*M Y-27632 ROCK Inhibitor for the first day after passaging. DMEM supplemented with 10% FBS and 1% Antibiotic antimycotic for BxPC3s. hiPSCs were harvested after 3–5 days of culture at 60–80% confluency, and differentiating cells were harvested on the stated day.

### 4.3 Flow cytometry

hiPSCs or hiPSC-NPCs were trypsinized for 5–7 minutes at 37 C and then, fixed with 4% v/v paraformaldehyde in PBS for 10-15 minutes. Cells were then permeabilized with 0.1% w/w Saponin in PBS for 5 minutes. Staining was performed with a 1:250 dilution of primary antibody and fluorescently conjugated isotype control antibody in 10% FBS in PBS blocking buffer overnight in the dark at 4C. Secondary fluorescent antibody was incubated for 1hr at room temperature. For data in which flow cytometry was performed directly after GEDI, cells were stained in primary antibody for 1 hour, then washed. Analysis was performed using a BD Accuri C6 flow cytometer (BD Biosciences, San Jose, CA) and Flowjo software. Cells were gated using FlowJo v10 first by identifying cells based on SSC:H vs FSC:H. Single cells were then identified by plotting FSC:A vs FSC:H and gating out cells that did not fit the main linear trend of this plot. Three biological replicates of undifferentiated hiPSCs were then fit to a normal distribution, and a cutoff threshold for counting cells as SSEA-5+ was set at just above the bottom 5% of undifferentiated hiPSCs.

### 4.4 Capture Experiments

Cells were labeled with 10*μ*M CellTracker Dye, either Green CMFDA or Red CMPTX (Thermo Fisher Scientific, Rockford, IL) for 45 minutes, trypsinized for 5–7 minutes at 37°C and resuspended in carrier solution (2% BSA, 1.3–1.5 mM EDTA in PBS) at approximately 300 cells/mL. Precise counts for each set of experiments were obtained by counting by hand 200ul samples of each cell-suspension-running-buffer mixture. In experiments in which cells were spiked into BxPC3 populations, cultures were trypsinized for 5–7 minutes at 37°C and resuspended in carrier solution at approximately 300,000 cells/mL. Functionalized silicon devices were mounted with Tygon tubing inlets and outlets in a custom PMMA holder. Cell capture was achieved by flowing 0.5 mL of cell suspension through the device at 1 mL/h followed by manual cell enumeration using fluorescence microscopy.

### 4.5 On-Chip Immunofluorescence

Cells were trypsinized for 5–7 minutes at 37°C and resuspended in carrier solution (2% BSA, 1.3–1.5 mM EDTA in PBS) at approximately 800,000-1,000,000 cells/mL. 0.5 mL of cell suspension was processed through GEDI chip functionalized with anti-SSEA-5 or normal mouse IgG (control) antibodies. Samples were fixed in 2% PFA in 50% PHEM buffer (60 mM PIPES, 25 mM HEPES, 10 mM EGTA and 2 mM MgCl2) for 15 minutes and blocked in 10% FBS in PBS for 1 hour. After staining of surface markers, the samples were permeabilized with 0.1% (w/w) Saponin. The samples were stained with, for Oct4, a primary (Abcam, ab60720,) and an AlexaFluor-488-conjugated secondary antibody (Invitrogen), for SSEA-3, a primary Rat monoclonal to SSEA-3 (Abcam, Ab16286, Lot 1987272) and an AlexaFluor-555-conjugated secondary goat anti-rat antibody (Abcam, A21434, Lot GR3225471-1) and, for DNA/nuclei, DAPI (Invitrogen). Samples were processed, fixed and stained within 48 h of collection.

### 4.6 GEDI Effluent ImmunoFluorescence

Three 60mm plates (Corning, CLS430166) of Day 9 hiPSC-NPCs were mixed and pumped through anti-SSEA-5 and isotype control functionalized GEDI chips at 1mL/hr for 30 minutes. The effluent of each chip (3 of each functionalization) was collected and immeditately fixed for 12 minutes in 4% w/v PFA. 70% of cells were then stained for Oct4, Nestin, and SSEA-3. 30% of cells received the corresponding secondary fluorescent antibody only. After staining, cells were pipetted and mounted onto glass coverslips coated in alginate hydrogel, to stick the fixed cells to the glass. During mounting, 2 of the secondary control slides (1 SSEA-5 and 1 isotype conrol) were damaged. Cells were then located on slides via the DAPI signal. Each slide was imaged in 3-4 locations on a Zeiss Z710 confocal microscope.

#### 4.6.1 CellProfiler Image Analysis

Quantitative data was obtained by using DAPI signal to locate cells as primary objects in CellProfiler. Secondary objects were then identified using DAPI as seed locations for secondary objects. The total intensity of the secondary object (ie: Oct4 or SSEA-3 intensity) was calculated. Mean Oct4 and SSEA-3 intensity across each of the four conditions (anti-SSEA-5/positive stain, anti-SSEA-5/control stain, isotype control/positive stain, and isotype control/negative stain) for all objects was calculated, and divided by mean DAPI object area for the same condition. Secondary object size was normalized by DAPI area, due to the fact that secondary object area had high variance in the negative stain, due to poorly defined object borders (as expected). Mean secondary object positive stain fluorescence intensity normalized by mean secondary object negative stain fluorescence intensity is reported in **Figure 4**. **Figure 4** also reports the fraction of secondary objects in the positive stain that are a specified fraction brighter than the mean negative stain corresponding secondary object (ie: a threshold of 1.5 means the object is counted as positive if its integrated intensity is at least twice 1.5 times as bright as the average corresponding negative stain object.

### 4.7 Statistics

All statistical tests were performed in GraphPad 8.0 using either a one-way ANOVA test using either Tukey’s multiple comparison test or student’s t test, or Dunnett’s multiple comparison test, where specified. Box and whiskers plots show all points, min to max, middle bar represents mean.

## Acknowledgements

The authors would like to acknowledge funding support from the National Science Foundation award number 1511194, awarded to B.K. and A.S., and partial support from NSF CAREER award 1554275 (A.S.) The authors would like to acknowledge Tyler Kirby and Jan Lammerding for supplying the hiPSC cells and for providing training in the culture of these cells.

## Author contributions statement

All experiments were performed by K.W. Pass through experiments were performed by K.W. and C.L. Data was analyzed by K.W., A.S., and B.K. Experiments were conceived by K.W., A.S., and B.K. K.W., A.S, and B.K. wrote the paper.

