## Supplementary figures and images for "Microfluidic Chip for Label-Free Removal of Teratoma-forming Cells from Therapeutic Human Stem Cells"

### Supplementary Figure 1

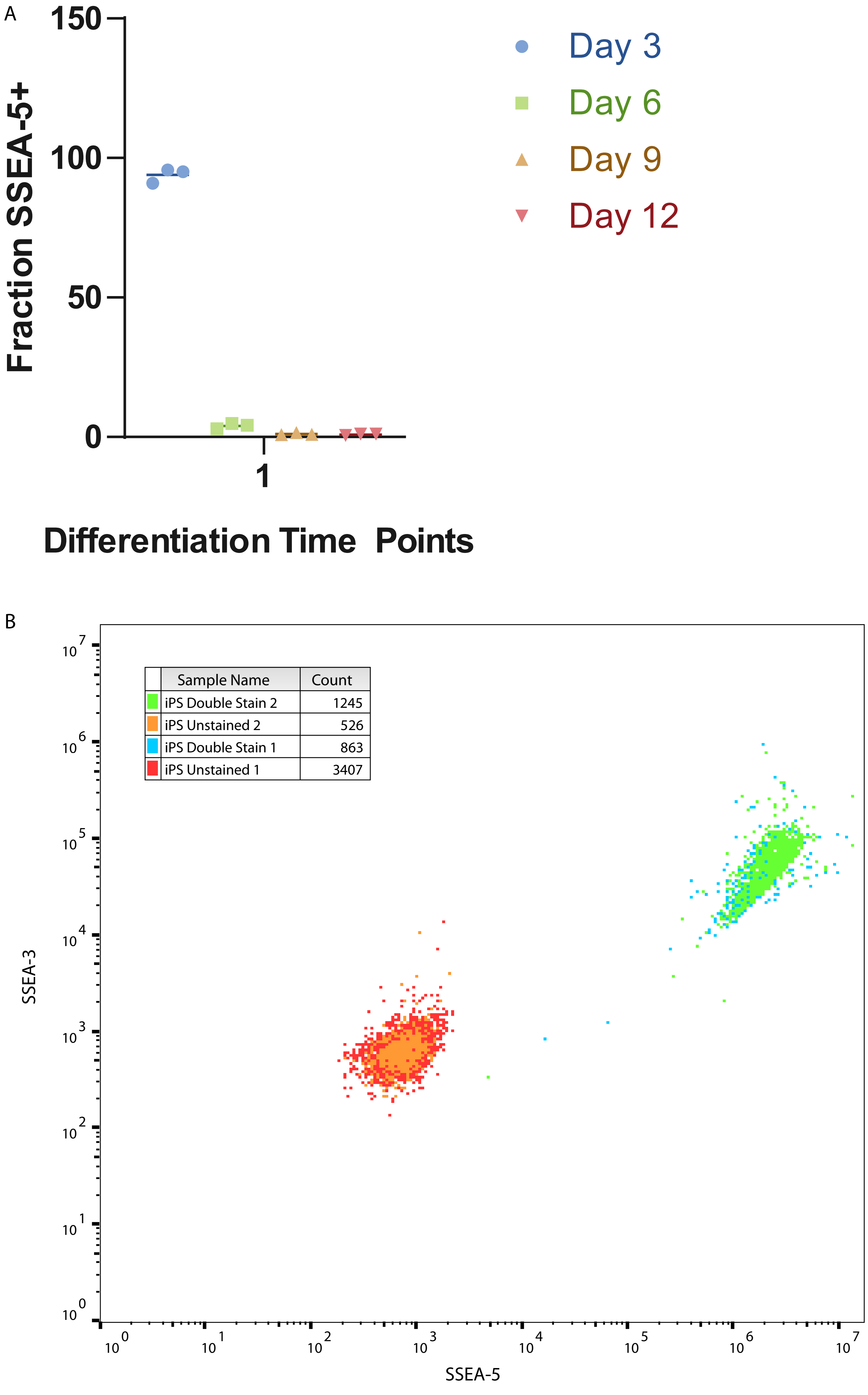
